# A Trizol-Beadbeater RNA extraction method for yeast, and comparison to Novogene RNA extraction

**DOI:** 10.1101/2025.07.30.667760

**Authors:** S. Honey, B. Futcher

**Affiliations:** Dept. of Microbiology and Immunology, Stony Brook University, Stony Brook, NY

## Abstract

58 samples of *S. cerevisiae* were sent to the Novogene Corporation for the intended purpose of RNA extraction, cDNA preparation, and sequencing. RNA was extracted by Novogene. However, yields of total RNA were about 1%, perhaps 1.25%, of the theoretically expected amount, which seemed low. Therefore, we extracted RNA locally using Trizol and bead-beating from a second, identical aliquot of 55 of the samples sent to Novogene. This local extraction gave yields that were on average 40% of the theoretical expectation, about 40 fold higher than the yields obtained by Novogene.

## Introduction

RNA-Seq is a standard method of molecular biology (Wang, Gerstein, & Snyder, 2009). For organisms with small genomes such as *S. cerevisiae*, it is cost-effective to pool many sequencing libraries on one flow cell. However, it is possible for a badly-prepared sequencing library to interfere with the sequencing of other libraries on the same flow cell. Therefore, commercial sequencing enterprises often insist that they make the cDNA sequencing libraries in-house for shared flow-cells, often starting at the step of RNA extraction from cellular samples. The ability of these commercial enterprises to extract RNA from various types of biological samples could affect results.

The difficulty and efficiency of RNA extraction varies with the nature of the cellular samples. Isolated mammalian cells are bounded by only a lipid membrane, and are relatively easy to lyse. In contrast, fungi such as the yeast *S. cerevisiae* have a tough cell wall (as well as a membrane), and it is generally necessary to break the cell wall, or permeabilize it in some way, to extract RNA (Collart & Oliviero, 2001; DeCaprio & Kohl, 2020; Green & Sambrook, 2021; Sasidharan, Amariei, Tomita, & Murray, 2012; Schmitt, Brown, & Trumpower, 1990). This can be difficult, and can require special equipment, precise methods, and expertise.

Here we compare the efficiency of RNA extraction from yeast at Novogene by their methods to the efficiency in our own laboratory by a Trizol-bead-beating method.

## Results

We had an RNA-Seq project requiring sequencing of 58 yeast RNA-Seq libraries. Although we are generally successful at extracting RNA from yeast and making sequencing libraries, most sequencing vendors prefer to extract the RNA and make sequencing libraries themselves. However, extracting RNA from *S. cerevisiae* can be challenging. We contacted multiple vendors, and asked about their experience and expertise in extracting RNA from yeast. While some vendors said they had little such experience, Novogene replied that they had extensive experience and expertise. Therefore we contracted with Novogene for our RNA-Seq project.

We collected fifty-eight 50 ml samples of yeast cells of defined size by elutriation. Concentrations ranged from 0.21 to 2.2 × 10^7 cells per ml, so 50 ml samples contained between 1.05 and 11 × 10^8 cells, with an average of 4.2 × 10^8. Since a yeast cell contains about 7.1 × 10^-13 g RNA per cell (von der Haar, 2008) (https://bionumbers.hms.harvard.edu/bionumber.aspx?s=n&v=4&id=104311), one expects an average of nearly 300 micrograms of total RNA per 50 ml sample. (However, since these cells were collected by elutriation, and were smaller than cells in asynchronous populations, we estimate that the average sample probably contained somewhat less than that).

Cells were collected by centrifugation and resuspended in 1 ml Trizol, per Novogene’s instructions. Initially, 0.5 ml of this 1 ml suspension was sent to Novogene for processing. (Subsequently, the other 0.5 ml of three samples (SH8, SH30, and SH55) was also sent to Novogene for processing.) Since 0.5 ml was half the total sample, the theoretical expectation for the average amount of total RNA was about 150 micrograms (or somewhat less, given the small average cell size).

After processing, Novogene reported an average recovery of only 0.71 micrograms of total RNA per sample, ranging from 0.03 micrograms to 2.63 micrograms (Table 1). This is only a small fraction of the ∼150 micrograms of total RNA expected.

**Table 1.**

| Sample | Cells<br>($\times 10^7$) | Theoret.<br>RNA ($\mu\text{g}$) | SB<br>RNA ($\mu\text{g}$) | Novogene<br>RNA ($\mu\text{g}$) | Novo/<br>Theoret. | Novo/<br>SB |
| --- | --- | --- | --- | --- | --- | --- |
| SH 1 | 34.5 | 245 | 28.6 | 0.32 | 0.0013 | 0.0112 |
| SH 2 | 23.3 | 165 | 41.3 | 0.29 | 0.0018 | 0.0070 |
| SH 3 | 27.5 | 195 | 38.9 | 0.27 | 0.0014 | 0.0069 |
| SH 4 | 31.5 | 224 | 71.6 | 0.62 | 0.0028 | 0.0087 |
| SH 5 | 9.0 | 64 | 17.2 | 0.76 | 0.0119 | 0.0442 |
| SH 6 | 9.0 | 64 | 26.7 | 0.74 | 0.0116 | 0.0277 |
| SH 7 | 7.8 | 55 | 14.1 | 0.53 | 0.0096 | 0.0377 |
| SH 8 | 7.8 | 55 | 4.6 | 0.03 | 0.0005 | NA |
| SH 9 | 8.0 | 57 | 18.0 | 0.87 | 0.0153 | 0.0482 |
| SH 10 | 8.0 | 57 | 19.6 | 0.97 | 0.0171 | 0.0494 |
| SH 11 | 7.8 | 55 | 14.1 | 1.46 | 0.0265 | 0.1036 |
| SH 12 | 7.8 | 55 | 33.6 | 1.16 | 0.0211 | 0.0345 |
| SH 13 | 15.3 | 108 | 38.2 | 0.20 | 0.0018 | 0.0052 |
| SH 14 | 15.3 | 108 | 34.4 | 0.20 | 0.0018 | 0.0058 |
| SH 15 | 11.0 | 78 | 45.8 | 0.49 | 0.0063 | 0.0107 |
| SH 16 | 11.8 | 83 | 73.1 | 0.40 | 0.0048 | 0.0055 |
| SH 17 | 11.3 | 80 | 48.0 | 1.20 | 0.0150 | 0.0250 |
| SH 18 | 11.3 | 80 | 40.7 | 0.66 | 0.0083 | 0.0162 |
| SH 19 | 12.5 | 89 | 56.5 | 0.82 | 0.0092 | 0.0145 |
| SH 20 | 12.5 | 89 | 94.2 | 2.27 | 0.0256 | 0.0241 |
| SH 21 | 55.0 | 391 | 63.2 | 0.56 | 0.0014 | 0.0089 |
| SH 22 | 55.0 | 391 | 44.1 | 0.62 | 0.0016 | 0.0141 |
| SH 23 | 50.0 | 355 | 79.6 | 1.17 | 0.0033 | 0.0147 |
| SH 24 | 52.5 | 373 | 91.2 | 1.14 | 0.0031 | 0.0125 |
| SH 25 | 50.0 | 355 | 132.3 | 0.69 | 0.0019 | 0.0052 |
| SH 26 | 50.0 | 355 | 128.6 | 0.88 | 0.0025 | 0.0068 |
| SH 27 | 50.0 | 355 | 110.0 | 0.95 | 0.0027 | 0.0086 |
| SH 28 | 50.0 | 355 | 122.5 | 1.55 | 0.0044 | 0.0127 |
| SH 29 | 13.3 | 94 | 14.0 | 0.25 | 0.0027 | 0.0178 |
| SH 30 | 11.3 | 80 | 3.8 | 0.19 | 0.0024 | NA |
| SH 31 | 11.8 | 84 | 56.5 | 0.22 | 0.0026 | 0.0039 |
| SH 32 | 11.0 | 78 | 56.2 | 0.22 | 0.0028 | 0.0039 |
| SH 33 | 11.0 | 78 | 43.7 | 0.28 | 0.0036 | 0.0064 |
| SH 34 | 11.5 | 82 | 32.8 | 0.43 | 0.0053 | 0.0131 |
| SH 35 | 11.5 | 82 | 96.3 | 1.22 | 0.0149 | 0.0127 |
| SH 36 | 19.5 | 138 | 59.8 | 0.32 | 0.0023 | 0.0053 |
| SH 37 | 19.5 | 138 | 26.2 | 0.45 | 0.0033 | 0.0172 |
| SH 38 | 17.8 | 126 | 52.2 | 0.26 | 0.0021 | 0.0050 |
| SH 39 | 18.0 | 128 | 53.0 | 0.41 | 0.0032 | 0.0077 |
| SH 40 | 17.5 | 124 | 70.0 | 0.77 | 0.0062 | 0.0110 |
| SH 41 | 18.0 | 128 | 39.3 | 0.72 | 0.0056 | 0.0183 |
| SH 42 | 18.0 | 128 | 57.9 | 1.07 | 0.0084 | 0.0185 |
| SH 43 | 18.0 | 128 | 56.3 | 1.72 | 0.0135 | 0.0306 |
| SH 44 | 30.0 | 213 | 50.3 | 0.68 | 0.0032 | 0.0135 |
| SH 45 | 30.0 | 213 | 40.2 | 0.63 | 0.0030 | 0.0157 |
| SH 46 | 25.0 | 178 | 84.9 | 0.63 | 0.0035 | 0.0074 |
| SH 47 | 25.0 | 178 | 80.6 | 0.86 | 0.0048 | 0.0107 |
| SH 48 | 30.0 | 213 | 76.7 | 0.79 | 0.0037 | 0.0103 |
| SH 49 | 27.5 | 195 | 92.6 | 1.22 | 0.0062 | 0.0132 |
| SH 50 | 27.5 | 195 | 74.7 | 2.63 | 0.0135 | 0.0352 |
| SH 51 | 7.5 | 53 | 9.7 | 0.59 | 0.0111 | 0.0610 |
| SH 52 | 7.5 | 53 | 6.9 | 0.38 | 0.0071 | 0.0551 |
| SH 53 | 5.3 | 38 | 15.3 | 0.25 | 0.0066 | 0.0164 |
| SH 54 | 5.3 | 38 | 16.9 | 0.3 | 0.0079 | 0.0178 |
| SH 55 | 5.5 | 39 | 1.8 | 0.25 | 0.0064 | NA |
| SH 56 | 6.0 | 43 | 23.0 | 0.5 | 0.0117 | 0.0217 |
| SH 57 | 5.5 | 39 | 23.2 | 0.43 | 0.0110 | 0.0185 |
| SH 58 | 5.5 | 39 | 17.1 | 0.44 | 0.0113 | 0.0257 |
| Mean |  | 143 | 49.4 | 0.71 | 0.007 | 0.0193 |

“Cells ( $\times 10^7$ )” is the number of cells in each sample.
“Theoret. RNA” is the theoretical amount of RNA in each sample, based on each cell containing $7.1 \times 10^{-7}$ micrograms of RNA. However, this is likely a slight to moderate over-estimate in most of these cases, since cells in most of these samples were smaller than usual, due to the nature of the experiment that produced them.
“SB RNA” is the number of micrograms of RNA produced locally (i.e., at Stony Brook) from these samples. For SH8, SH30, and SH55, the sample was the residual cell suspension, much less than a full 0.5 ml sample.
“Novogene RNA” is the number of micrograms of RNA produced by Novogene from these samples.
“Novo/Theoret.” is the ratio of the amount of RNA produced by Novogene to the theoretical amount of RNA.
“Novo/SB” is the ratio of the amount of RNA produced by Novogene to the amount of RNA produced at Stony Brook. For SH8, SH30, and SH55, this is “NA” because a full sample was not available at Stony Brook.

The yield of RNA seemed alarmingly low. In particular, we were concerned that the low yields might reflect a small proportion of lysed cells, and that aberrant cells might lyse more easily than average, and that the RNA recovered might therefore represent these aberrant cells, and not the average cell. In such a case, RNA-Seq results would be misleading, as they would represent gene expression in aberrant cells.

To investigate the low RNA yields further, we processed the other 0.5 ml of 55 of the original samples ourselves. (In fact, the volumes still available were somewhat less than 0.5 ml.) In addition, for samples SH8, SH30, and SH55, there was some residual cell suspension in the original tubes, and we recovered several microlitres of the residual material and extracted RNA. The samples had been stored at -80°C in Trizol, and we used a zirconium bead/bead-beater protocol to lyse the cells, still in Trizol (see Methods).

Results are shown in Table 1. Table 1 shows the number of cells in each sample, the theoretical amount of RNA in each sample, the amount of RNA recovered by Stony Brook or Novogene for each sample, the ratio of the amount of Novogene RNA to theoretical RNA; and the ratio of the amount of Novogene RNA to Stony Brook RNA. Since the sample volumes we were actually able to recover were less than 0.5 ml, Table 1 underestimates the yield of our method. Also, we are reporting the final yield of pure total RNA after all RNA recovery and processing steps.

We obtained an average of 49.4 micrograms of total RNA per sample, compared to 0.71 micrograms of total RNA per sample at Novogene. When the yields in micrograms were compared between each sample by a paired t-test, the difference was significant with a p-value of 2 × 10^−16^.

When we calculated yields for each sample, then averaged these 55 yields, we found that on average we obtained 39.4% of the theoretical yield, versus 0.70% for Novogene, a difference of about 50-fold. Even for the three samples where we had only residual material, we obtained more RNA than did Novogene.

We went on to use the RNAs prepared locally to make cDNA libraries, and to sequence these cDNA libraries (RNA-Seq) obtaining very large libraries and excellent sequencing results (Honey and Futcher, in preparation).

## Discussion

RNA extraction from *S. cerevisiae* was very inefficient at Novogene (about 1% efficiency) compared to our method (about 40% efficiency).

This low efficiency could be because only about 1% of cells in the sample were lysed. Although there is no proof that inefficient lysis is the cause, there are two supporting arguments. First, the method possibly used by Novogene to lyse the cells would be expected to be inefficient (see Methods). Second, the yeast samples where Novogene was more successful (∼ 3% efficiency) were samples that contained a relatively high proportion of large, aberrant cells, and the yeast samples where Novogene was less successful (∼ 0.05% efficiency) were samples containing mainly small, round cells. Alternatively, the low efficiency could have been due to inefficient RNA recovery (see Methods).

If cell breakage were inefficient but completely random, then the ensuing RNA-Seq results could still be useful. However, if cell breakage occurred preferentially for large, aberrant cells, then the results from RNA-Seq would represent gene expression in these aberrant cells, and so would be misleading with regard to gene expression in the entire sample population. Thus it could be potentially misleading to sequence RNA samples when extraction efficiency is very low.

It was difficult to find out from Novogene what methods they had used, and this is problematic in terms of identifying the reason for low yields, and generally for publishing a scientific report. Also, although Novogene has several quality controls in place at various steps of the RNA-Seq pipeline, it seems they do not have any quality control for RNA yield.

While we have found a low efficiency of Novogene RNA extraction for yeast, yeast is certainly a difficult case for RNA extraction. We have no reason to think that Novogene’s RNA extraction efficiency would be poor in easy cases, such as in mammalian tissue culture cells. However, in the apparent absence of any quality control for the extraction step, it seems prudent for investigators to check the RNA yield for any kind of sample. This is a parameter that should be reported in publications.

## Methods

### Local (Stony Brook) methods

#### Yeast cell lysis

Our method combines Trizol for solubilizing and stabilizing RNA with bead-beating to break the yeast cell wall. Trizol is a common reagent for extraction of RNA (Rio, Ares, Hannon, & Nilsen, 2010), and is a trade-name for an acidic mixture of guanidinium thiocyanate, phenol, and chloroform. It was originally developed by Chomczynski and Sacchi (Chomczynski & Sacchi, 1987, 2006). Bead-beating refers to the process of mixing yeast (in some buffer) with solid beads of material, which may be glass, or zirconium oxide, or carborundum (silicon oxide), and then mechanically agitating the yeast-bead suspension in some way. The beads are more effective if they are denser (zirconium oxide and carborundum are denser than glass), and if they are the right size (for yeast, usually 0.2 to 0.5 mm), and if the mechanism of agitation achieves high acceleration and velocity (a Bead Ruptor 12 can achieve velocities of 6 meters per second). Many protocols for extraction of RNA from yeast have been published (DeCaprio & Kohl, 2020; Green & Sambrook, 2021; Kruckeberg, Nagarajan, McInnerney, & Rosenzweig, 2009; Lee, Hong, & Kang, 2019; Rivas, Vizcaino, Buey, Mateos, Martinez-Molina, & Velazquez, 2001; Sasidharan et al., 2012; Shedlovskiy, Shcherbik, & Pestov, 2017), and our protocol uses many of the same methods as used by these and other previous workers.

Yeast suspended in Trizol were placed in a screw-cap (with O-ring) 2 ml tube. 0.5 mm zirconium oxide beads were poured from a plastic weigh-boat into the 2 ml tube, until the level of beads was about 1 mm below the meniscus of the liquid. Tubes were chilled to 0°C on ice, then were fastened (in a balanced fashion) into an Omni International Bead Ruptor 12. The Bead Ruptor speed (S) was set to 3.1 meters/sec, and the time (T) set to 30 sec. (Prolonged treatment should be avoided, as the sample heats rapidly during the intense agitation.) After one 30 second treatment, tubes were removed and placed in ice. Samples of 1 microlitre of liquid (avoiding beads) were removed and pipetted into 6 microlitres of water on a microscope slide. The cells on the slide were inspected using phase microscopy for breakage. Typically breakage was greater than 85%. In case of poor breakage, a second 30 second cycle would have been done, however no second cycle was needed for any of the 55 samples considered here.

(The Bead Ruptor uses a complex motion which we find highly effective. In the past, we have used other types of oscillating machines which have a back-and-forth motion along one dimension. For these types of machines, there can be a problem with certain sample volumes, such that the sample (beads plus yeast plus buffer) stays in essentially the same place, while the tube oscillates around it. In this rare case, there is little cell breakage. Adjusting the volume of the tube, the volume of the sample; the length of the oscillatory path; or sometimes the frequency of oscillation, can restore breakage.)

#### RNA recovery and processing

The cell lysate was transferred to a 1.5 mL Eppendorf tube using a pipette tip, leaving the zirconium oxide beads behind in the original 2 mL tube. Chloroform (20% of the volume of TRIzol used) was added to the lysate, and the contents were mixed by gently rocking for 3 minutes. The mixture was then centrifuged at 14,000 rpm in a tabletop centrifuge for 15 minutes. This step resulted in phase separation into three layers: a colorless upper aqueous phase containing partially purified RNA, an interphase containing DNA, and a lower red phenol–chloroform phase containing proteins. The upper aqueous phase was carefully transferred to a 15 mL tube, avoiding contamination from the interphase or lower layer.

To the recovered aqueous phase, 350 µL of binding buffer (from Ambion RiboPure™ RNA Purification Kit – Yeast, Cat #AM1926) was added per 100 µL of aqueous solution. RNA precipitation was carried out by adding 200-proof ethanol to achieve a final ethanol concentration of 70%. The mixture was then processed using the filter cartridges supplied with the Ambion kit, following the manufacturer’s protocol. RNA was eluted from the cartridges with 2 × 25 µL of 10 mM Tris-HCl buffer (pH 8.0). The RNA concentration was quantified using a NanoDrop spectrophotometer.

### Statistics

Stony Brook and Novogene yields (micrograms) were compared using a paired t-test (Excel).

Averages. Two approaches were used for averaging, as was most appropriate to the results being analyzed. One was to sum over all samples, then divide by the number of samples; the other was to calculate a result for each sample, then average these results. For instance, when we state in the text: “We obtained an average of 49.4 micrograms per sample, compared to 0.71 micrograms per sample at Novogene.”, this was the sum-then-divide method: the number of micrograms was summed over all samples, then divided by the number of samples. However, when we state in the text: “When we calculated yields for each sample, then averaged these 55 yields, we found that on average we obtained 39.4% of the theoretical yield, versus 0.70% for Novogene, a difference of about 50-fold.” this was the calculate-then-average approach: yields were calculated individually for each of the 55 samples, then the 55 yields were averaged. The two approaches give similar but not identical answers; it appears the answers are not identical mainly because Novogene yields were especially bad for the samples with the most RNA (e.g., SH21 through SH28), and this skews one average versus the other.

### Cell lysis (Novogene)

We spoke to several customer service representatives at Novogene to try and find out what methods had been used for cell breakage, but we were not able to speak to laboratory personnel. We obtained several different answers about methods, but the most specific answer was:

“Our team used mechanical disruption using type C bead tubes from MN and a TissueLyser II by Qiagen operated at 30 Htz for 5 minutes per side. There was no mention of an assay for cell breakage.”

If this is the method that was used, one problem is the rate of oscillation, 30 Htz (i.e., 30 cycles of oscillation every second). The throw-distance for the oscillation of a TissueLyser II is roughly 2 cm (perhaps less). 30 cycles of oscillation every sec means a speed of roughly 120 cm/sec (assuming a 2 cm throw), whereas the Bead Ruptor was set to 3100 cm/sec, about 25 times faster. Furthermore kinetic energy is proportional to the square of velocity, so the energy developed in the Bead Ruptor may have been 625-fold higher. The absence of an assay for breakage is also problematic.

#### RNA Recovery (Novogene)

A customer service representative thought RNA may have been recovered using the RNeasy Mini kit-Trizol, which would have selectively recovered RNAs over 200 bp in length. 15% to 20% of yeast RNA (mainly tRNA, 5S RNA, 5.8S RNA) is less than 200 bp. If this was the method used at Novogene, then their yields should be considered about 25% relatively higher than reported above. That is, their yields would have been roughly in the range of 0.875% to 1.25%. It is possible that the relatively poor yields at Novogene reflect inefficient RNA recovery, rather than inefficient cell breakage. In this case, the low yields might not be specific to yeast.

